# LFA-1 Interaction with GBP-130 on *Plasmodium falciparum*-infected Red Blood Cells mediates NK Cell Activation and Parasite Control

**DOI:** 10.1101/2025.08.20.671208

**Authors:** Osama Mukhtar, Ravi Dutt, Ashutosh Panda, Poonam Kumari, Suneet Shekhar Singh, Gourab Paul, Neha Prakash, Madiha Abbas, Md. Muzahidul Islam, Priya Arora, Alma Tammour, Asif Mohmmed, Dhiraj Kumar, Pawan Malhotra

## Abstract

Natural Killer (NK) cells contribute to early immunity against *Plasmodium falciparum* by recognizing and eliminating infected red blood cells (iRBCs), a process mediated in part by the integrin LFA-1. However, the cognate parasite ligand for LFA-1 has remained unknown. Here, we identify Glycophorin Binding Protein-130 (*Pf*GBP-130) as a surface-expressed ligand on iRBCs that binds the I-domain of LFA-1 (LFA-1 αI). Using an LFA-1 αI-Fc fusion protein, we demonstrate stage-specific binding to iRBCs, and LC-MS/MS analysis of immunopreciptates of αI-Fc bound to iRBC revealed *Pf*GBP-130 as a high-confidence interactor. Recombinant *Pf*GBP-130 binds NK and THP-1 cells in an LFA-1-dependent manner. Co-culture assays show that *Pf*GBP-130 promotes NK cell activation, degranulation, and facilitates contact-dependent killing of iRBCs. Neutralizing antibodies against *Pf*GBP-130 significantly impair these responses. Our findings establish *Pf*GBP-130 as the LFA-1 ligand on iRBCs, providing new insight into NK cell-mediated immunity in malaria and identifying a potential target for host-directed interventions.

## Introduction

NK cells are innate immune cells involved in early defence against microbial pathogens, parasitic infections and tumor cells (Burrack et al., 2019). NK cells are found in various tissues including liver, peritoneal cavity as well as placenta and constitute up to 15% of peripheral blood lymphocytes. NK cells were initially identified due to their ability to kill certain tumor cells *in vitro* (Cerwenka and Lanier, 2001). Subsequent studies using *in vivo* murine models demonstrated that depletion of NK cells leads to enhanced tumor formation (Karre, 2002). Activation of NK cells results from the concerted action/response of cytokine signalling, adhesion molecules and the interaction of activating receptors with their corresponding ligands expressed on the surface of pathogenic infected cells or tumors. Despite having understood the role(s) of NK cells in controlling the microbial infection and tumor surveillance, mechanisms of NK cells recognition by pathogenic cells are not well understood.

Malaria infection and pathogenesis involve a complex series of interplays between parasite and human host factors and a better understanding of host cells (factors) may be important in developing innovative host derived therapeutic approaches. NK cells provide first line of defence against malaria parasite infection. Studies in mouse malaria models have shown that various immune cells such as natural killer (NK) cells, dendritic cells, T cells and B cells contribute to antiparasitic immunity (Chen et al., 2014; Ye et al., 2018). *Plasmodium* development in humans occurs in two phases: the merozoite stage in the liver and the symptomatic blood stage. During the liver stage, immunity against *Plasmodium* is primarily mediated by IFN-gamma. NK cells secrete IFN-gamma, which eliminates infected hepatocytes either directly or by activating other immune cells, including cytotoxic T cells and macrophages (Artavanis-Tsakonas and Riley, 2002; Burrack et al., 2019).

NK cells contribute to blood stage immunity through three main mechanisms. First, they facilitate the clearance of infected red blood cells (iRBCs) via a bystander mechanism, involving the secretion of IL-12 and IL-18. These cytokines prompt macrophages and dendritic cells to clear iRBCs or stimulate adaptive immune cells. Second, NK cells engage in antibody-dependent cell-mediated cytotoxicity (ADCC), wherein antibody-opsonized iRBCs are recognized via the CD16 (FcγRIII) receptor. This leads to the release of cytotoxic granules, including perforin and granulysin, enabling targeted lysis of infected cells. Third, NK cells can directly kill infected or transformed cells through activating receptor-ligand interactions, a mechanism well characterized in antiviral and tumor immunity. NK cells are critical in limiting acute malaria infection as depletion of NK cells in mouse malaria model has been associated with higher parasitemia and accelerated disease progression (Chen et al., 2014; Ye et al., 2018). NK cell protective effects are mediated through both cytotoxic activity and secretion of interferon-gamma (IFN-γ). Clinical studies in *P. falciparum*-infected children have revealed an inverse correlation between NK cell frequency or functional activity and parasite burden (Ojo-Amaize et al., 1981). Furthermore, experimental infections in malaria-naïve individuals have demonstrated that NK cells are among the earliest responders, capable of directly lysing iRBCs and releasing IFN-γ and soluble granzyme (Hermsen et al., 2003; Ye et al., 2018).

*Plasmodium* Infected RBCs (iRBCs) activates NK cell, enhancing IFN-γ production and cytotoxicity through perforin and granzyme release. This activation is contact-dependent, as demonstrated by the lack of upregulation of NK cell activation markers during transwell incubation. Co-incubation of iRBCs with NK cells promote *Plasmodium* parasite killing, and this contact-dependent interaction is mediated in part by the lymphocyte function-associated antigen-1 (LFA-1) on NK cells (Chen et al., 2014; Korbel et al., 2005). In this study, we demonstrate that iRBCs bind specifically to the “αI-domain” of LFA-1 and identified *Plasmodium* Glycophorin Binding Protein-130 (*Pf*GBP-130) as a major iRBC surface ligand interacting with LFA-1. Furthermore, we show that *Pf*GBP-130 expressed on the iRBC surface binds to THP-1 and NK cells, inducing activation and degranulation of these cells. Together, our findings identify *Pf*GBP-130 as a ligand for LFA-1 that facilitates direct contact-mediated killing of *Plasmodium*-infected erythrocytes by NK cells, highlighting its potential role in host–parasite immune interactions.

## Results

### LFA **α**I domain of Lymphocyte Function Associated Antigen-1 (LFA-1) is involved in ligand binding on iRBCs surface

Contact-dependent killing of *Plasmodium*-infected red blood cells (iRBCs) by NK cells is essential for effective malaria control and is mediated by LFA-1 (CD11a/CD18), a β2 integrin composed of an αL (CD11a) and β (CD18) subunit. Notably, CD11a contains an inserted (I) domain of approximately 200 amino acids, homologous to the von Willebrand factor type A (vWA) domain family (Suppl. Fig. 1), which plays a critical role in ligand binding, including interaction with ICAM-1(Qu and Leahy, 1995). Here, we generated an LFA-1 αI-Fc fusion protein to assess αI domain binding to iRBCs. Schematics of CD11a (a subunit of LFA-1) and it’s I domain fused to human IgG1 Fc domain is shown in Figure 1A. DNA sequences corresponding to LFA-1 αI-Fc fusion protein were cloned in a plasmid pFUSE-hIgG1-Fc2 (Suppl. Fig. 2A & 2B) and the protein was expressed in CHO K1 cells under serum-free conditions, Fusion protein was purified via protein-A affinity beads and purified protein was analysed by SDS-PAGE and western blot using secondary anti-human IgG. A protein of a ∼45 kDa was seen in SDS-PAGE and western blot (Fig. 1B), which corresponded to the molecular weight of fusion construct. We next analysed the binding of purified LFA αI-Fc protein with iRBCs and detected binding using secondary human antibody conjugated to PE Texas red or anti-human antibody conjugated to FITC. Binding was also analyzed for all the three asexual blood stages of *Plasmodium* ring, trophozoites and schizonts by FACS. As shown in figure 1C, high affinity binding was observed with all the three stages that was significantly higher than that of observed with the human IgG (hIgG) isotype control.

**Figure 1.**
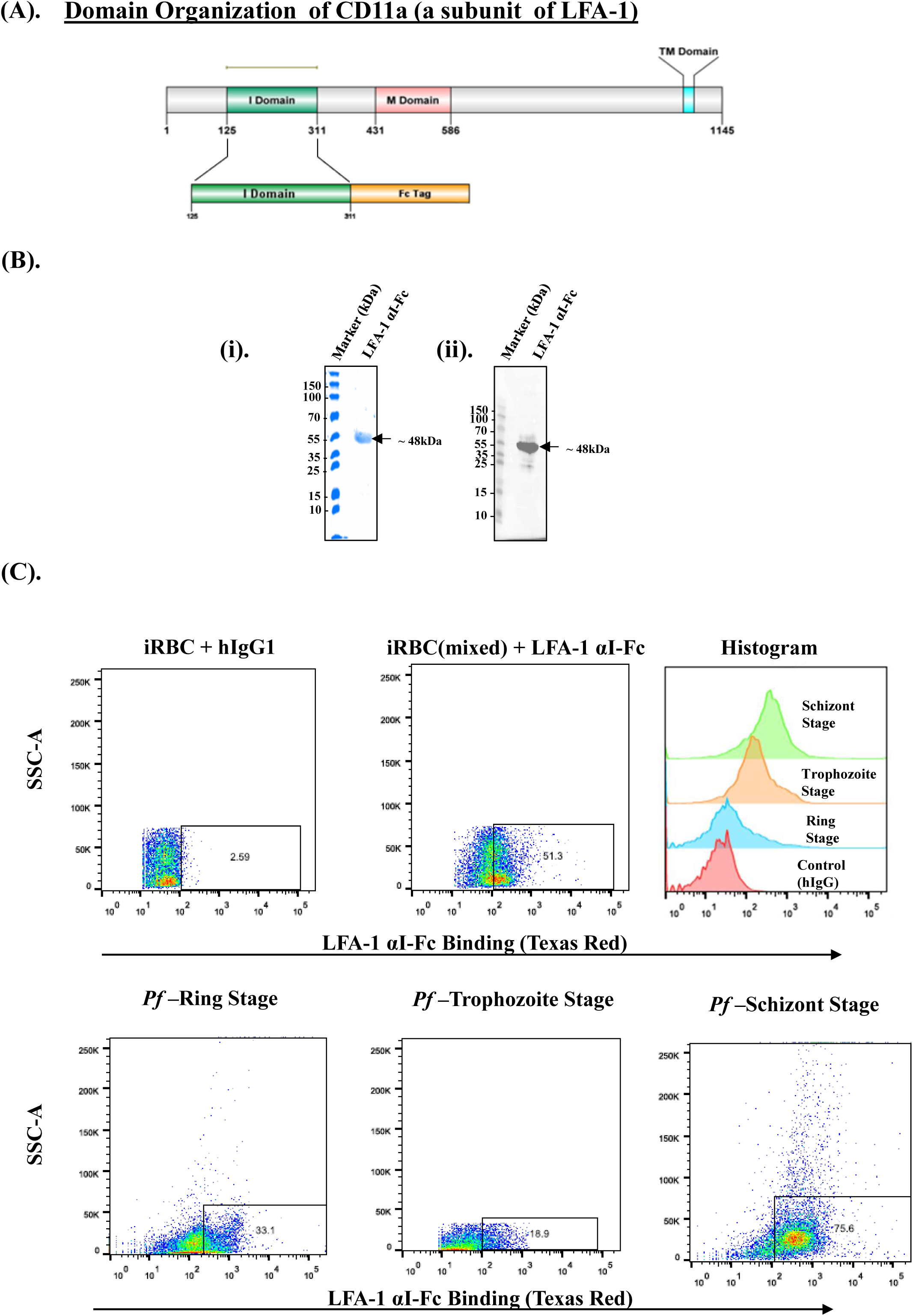
The. α**I domain of Lymphocyte Function-Associated Antigen-1 (LFA-1) binds to *Plasmodium falciparum*–infected erythrocytes.** (**A)** Schematic representation of full-length of CD11a, a subunit of LFA-1 and the recombinant construct encoding the C-terminal Fc-tagged αI domain of LFA-1. **(B)** Expression and purification of recombinant LFA-1 αI-Fc fusion protein. The αI domain of LFA-1 was cloned into the pFUSE-hIgG1-Fc2 vector and expressed in CHO-K1 cells. The fusion protein was purified from culture supernatants using Protein-A affinity chromatography. **(i)** SDS-PAGE analysis of the purified protein revealed a prominent Coomassie-stained band at ∼45 kDa, corresponding to the expected molecular weight of the LFA-1 αI-Fc fusion protein. **(ii)** Western blot analysis using anti-human IgG antibody confirmed the identity of the fusion protein. **(C)** Binding of LFA-1 αI-Fc (250nM) fusion protein to *P. falciparum*–infected red blood cells. Flow cytometry analysis using PE-Texas Red–conjugated anti-human IgG antibody demonstrated specific binding of the LFA-1 αI-Fc protein to ring, trophozoite, and schizont stages of iRBCs, indicating interaction across all major asexual blood stages.

To exclude the possibility that LFA-1 αI-Fc binding to infected red blood cells (iRBCs) resulted from non-specific interactions with erythrocyte surfaces rather than parasite-derived antigens, we performed parallel incubations using uninfected RBCs and an isotype-matched human IgG control. little or no binding was observed for these molecules to the infected cells. No significant difference in binding to uninfected RBCs was observed between LFA-1 αI-Fc and the isotype control, indicating the absence of non-specific erythrocyte binding (Suppl. Figure 2C). These results showed that the αI domain of LFA-1 is involved in binding to infected RBC cells.

### *P. falciparum* Glycophorin Binding Protein-130 protein on iRBC surface binds to LFA **α**I domain of LFA-1 protein

To elucidate the molecular basis of LFA-1 mediated interactions with *Plasmodium falciparum* infected erythrocytes, we undertook a comprehensive proteomic approach. Briefly, recombinant LFA-1 αI-domain Fc fusion protein (LFA-1 αI-Fc) used as an affinity reagent to capture iRBC surface-interacting proteins, following DTSSP cross-linking. Human IgG1 (hIgG1) served as a control to distinguish specific from non-specific background interactions. Immunoprecipitated proteins from both LFA-1 αI-Fc and hIgG1 conditions were analyzed by LC-MS/MS, enabling a comparative assessment of specifically interacting proteins. Three independent biological replicates were performed, and stringent selection criteria were applied such that only proteins supported by detection of ≥5 peptides were considered for downstream analysis. Although several proteins showed similar sequence coverage, these were also present in beads + human IgG control eluates, indicating non-specific background binding (Suppl. Table 1). Only proteins consistently detected across all replicates were considered. *Pf*GBP-130 (PF3D7_1016300) was identified as a top candidate based on peptide coverage and spectral abundance across all the replicates (Fig. 2A). LC-MS/MS analysis thus strongly suggested a specific and robust interaction between LFA-1 αI domain and *Pf*GBP-130.

**Figure 2.**
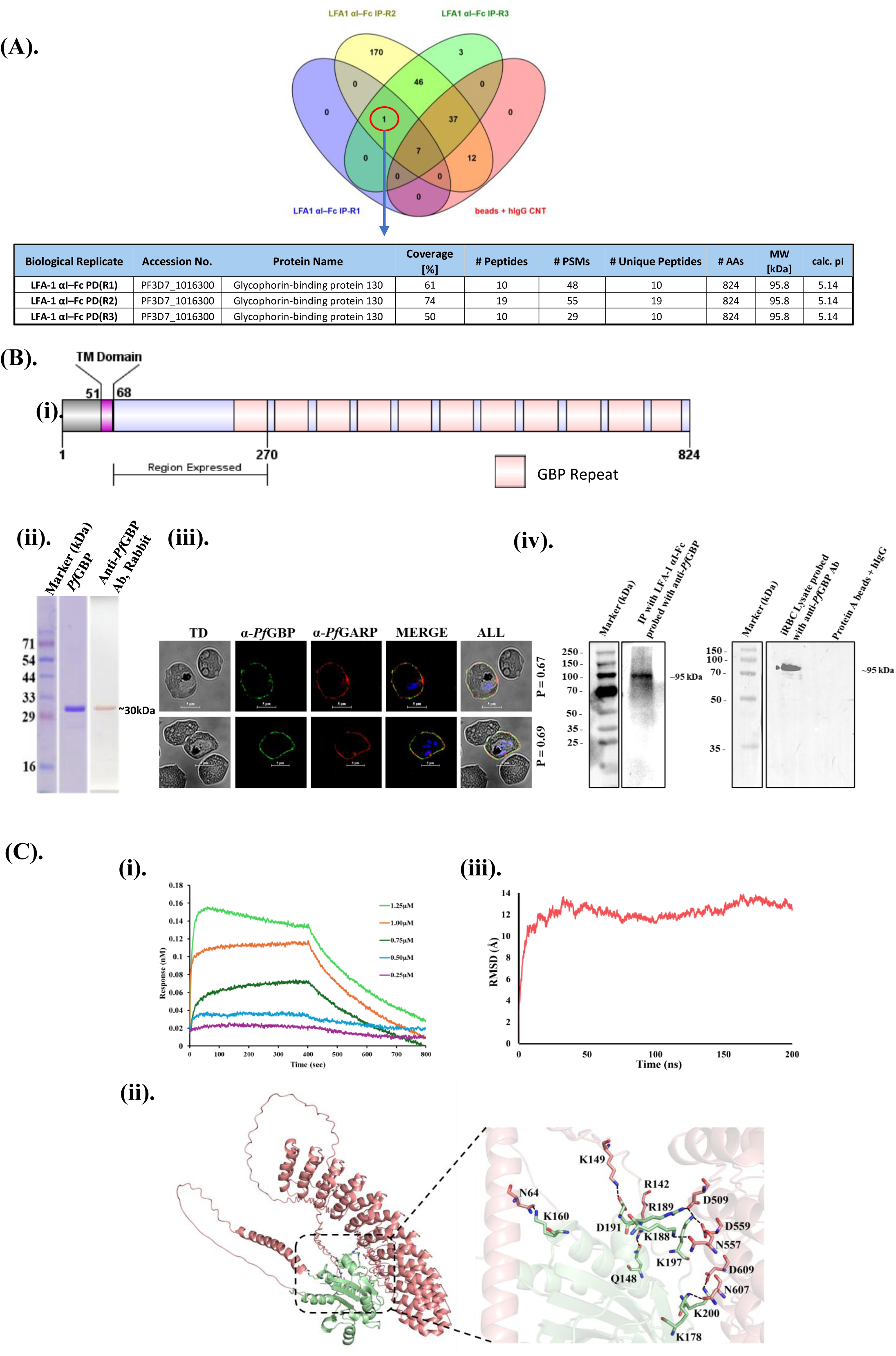
*Pf*GBP-130 on the surface of *Plasmodium falciparum*–infected erythrocytes bind the LFA-1 α**I domain**. **(A)** Identification of *Pf*GBP-130 as an interacting partner of LFA-1 αI-Fc. LC-MS/MS analysis of immunoprecipitates from *P. falciparum*–infected erythrocyte lysates pulled down with LFA-1 αI-Fc fusion protein revealed *Pf*GBP-130 (PF3D7_1016300) as a major interacting protein. The table summarizes proteins specifically enriched in the LFA-1 αI-Fc pull-down in all the three biological replicates, showing high peptide coverage and spectral abundance, indicating a specific and robust interaction. **(B)** Characterization and localization of *Pf*GBP-130. **(i)** Schematic representation of the domain organization of *Pf*GBP-130 and the N-terminal fragment (amino acids 69–270) that was expressed in *E. coli* (termed *Pf*GBP-130-N). **(ii)** SDS-PAGE and western blot analysis of purified *Pf*GBP-130-N using anti-rabbit *Pf*GBP-130 antibodies. A prominent band at ∼30 kDa corresponds to the expected molecular weight of the recombinant fragment. **(iii)** Immunofluorescence assay (IFA) demonstrating surface localization of *Pf*GBP-130 on trophozoite-stage iRBCs using anti-*Pf*GBP-130 antibodies. *Pf*GBP-130 (green) partially colocalizes with *Pf*GARP (red), a well-established iRBC surface protein with an extracellular domain. Nuclei were stained with DAPI (blue), confirming surface expression. **(iv)** Western blot analysis of iRBC lysate and of immunoprecipitate of LFA1 αI-Fc bound iRBC (figure 2A) using anti-rabbit GBP-130 antibody. Lane1, shows the presence of *Pf*GBP-130 in the immunoprecipitate. The ∼130kDa band is notably absent in the control IP eluate where no LFA-1 αI-Fc, was bound to iRBC, demonstrating the specificity of the LFA-1 αI-Fc and *Pf*GBP-130 interaction. Lane 2 shows the detection of native *Pf*GBP-130 as a ∼110 kDa protein in trophozoite stage *P. falciparum* lysate, consistent with its predicted molecular weight. **(C)** Biophysical and computational validation of *Pf*GBP-130 and LFA-1 αI interaction. **(i)** Bio-layer interferometry (BLI) analysis of real-time binding between *Pf*GBP-130-N and LFA-1 αI-domain. Averaged sensorgrams across independent experiments (n ≥ 3) demonstrate concentration-dependent association and dissociation kinetics. The calculated equilibrium dissociation constant (K_D_) was (1.7 ± 0.22) × 10[[M, indicating high-affinity binding. (**ii)** *In silico* docking analysis showing the energy-minimized complex of *Pf*GBP-130 (salmon) and LFA-1 αI domain (green) generated using ClusPro 2.0. Representative hydrogen bonds and interacting residues are shown as sticks. Visualizations were prepared using PyMOL. **(iii)** Molecular dynamics (MD) simulation of the *Pf*GBP-130/LFA-1 αI complex. The graph depicts the root mean square deviation (RMSD) over time, confirming structural stability of the protein–protein complex.

To validate the observed interaction and provide orthogonal confirmation of our proteomic findings, we expressed the N-terminal region of *Pf*GBP-130 (aa 69-270 encompassing one GBP repeat) in *Escherichia coli* and purified the protein to near homogeneity (Fig. 2B. i). Polyclonal antibodies were raised against recombinant GBP-130 N terminal protein fragment referred here as *Pf*GBP130-N in rabbit (Fig. 2B. ii). Antibodies raised against *Pf*GBP130-N were specific as they could detect the native *Pf*GBP-130 on iRBCs surface. We carried out co-localization IFA with *Pf*GARP, a well-established iRBC surface protein with an extracellular domain (Raj et al., 2020). Anti-*Pf*GBP130 N staining co-localized with *Pf*GARP at the iRBC surface, with a Pearson’s correlation coefficient of 0.67, supporting surface exposure of *Pf*GBP130 (Fig. 2B. iii). To confirm the specificity of anti-*Pf*GBP-130 N antibodies, we next performed a western blot analysis of trophozoite stage parasite lysate and also for the immunoprecipitaed fraction of LFA-1 αI-Fc protein bound iRBC lysate using anti-*Pf*GBP-130 N antibodies. As shown in figure 2B. iv, a distinct band at ∼95kDa, consistent with the reported molecular weight of *Pf*GBP-130 (Kochan et al., 1986), were observed exclusively within the LFA-1 αI-Fc bound iRBCs eluate as well as in iRBCs lysate control. Critically, this band was absent in the hIgG1 control eluate, confirming the specificity anti-*Pf*GBP-130 antibodies as well as of the observed interaction between *Pf*GBP-130 and recombinant LFA-1 αI-Fc protein, thus ruling out non-specific binding artifacts.

### Biophysical assessment of the interaction between *Pf*GBP130-N and LFA-1 **α**I-Fc fragments

To confirm the interaction between LFA-1 and PfGBP-130, real-time binding studies were performed using recombinant PfGBP-130N and the LFA-1 αI-domain by bio-layer interferometry (BLI) on the Sartorius Octet K2 system. Briefly, the LFA-1 αI-domain was immobilized on an AR2G sensor (Octet Amine Reactive Second-Generation biosensor) at 100 ng, and increasing concentrations of *Pf*GBP-130N were applied as analyte. *Pf*GBP-130N displayed a rapid association phase with immobilized LFA-1 αI-domain, followed by a clear dissociation phase, consistent with direct binding. To ensure robustness and reproducibility, the BLI experiments were performed in multiple independent replicates (n ≥ 3) using independently purified protein batches. The averaged sensorgrams, together with the standard deviation of the calculated equilibrium dissociation constant, yielded a K_D_ of (1.7 ± 0.22) × 10 M (Figure 2C(i)), indicating a strong and specific interaction between LFA-1 αI-domain and *Pf*GBP-130N.

Since LFA-1 αI-domain displayed a high binding affinity for *Pf*GBP-130 N, we next performed protein-protein docking studies to examine the molecular-level interactions between these two proteins. The *Pf*GBP-130 model was docked against LFA-1 αI-domain using the Cluspro2.0 protein-protein docking server, and the binding site for the generated complex was analyzed with PyMOL. The best resulting structure was submitted to PDBSum to identify the amino acid residues involved in the interactions. Examination of the interaction surface showed that LFA1 binds within a curved pocket of *Pf*GBP130 formed by an anti-parallel helix structure, and the interacting residues for LFA1 were mainly localized in the N-terminal domain. A total of 13 hydrogen bonds and five salt bridges were observed for the *Pf*GBP130/LFA1 complex. The representative hydrogen bonds are shown in figure 2C. ii.

Furthermore, the binding energy of the docked LFA1/*Pf*GBP complex was evaluated utilizing several computational tools, including PRODIGY, PPCheck, AREA-AFFINITY, and the HawkDock server, each of which consider different parameters to calculate the energy. As summarized in Suppl. Table 2, the calculated energy values suggest stable complex formation.

To assess the strength of the interactions, the buried surface area of the complex was calculated using the PDBePISA web server, yielding a value of 1528.7 Å² per molecule, which is in accordance with the buried area suggested for known protein-protein complexes. Next, we evaluated the stability of the docked complex through molecular dynamics (MD) simulations in an aqueous environment over 200 nanoseconds (ns). The Root Mean Square Deviation (RMSD) trajectory plot, indicated that after a minor fluctuation during the first 30 ns, the complex remained stable for the rest of the simulation (Fig 2C. iii). In conclusion, both real-time BLI and *in-silico* studies suggested a stable interaction between LFA-1 and *Pf*GBP-130.

### *Pf*GBP-130 ectodomain shows a specific LFA-1-dependent binding to human THP-1 monocytic and primary NK cells

*Pf*GBP-130 possesses an N-terminal cytosolic domain, a transmembrane domain, and an extracellular domain characterized by repeats (Fig. 3A). Since, we identified *Pf*GBP-130 as a putative ligand for LFA-1 on NK cells, we next investigated the direct interaction of *Pf*GBP-130 with immune cells, specifically THP-1 monocytes and NK cells. Briefly, recombinant protein construct comprising of the extracellular domain of *Pf*GBP-130, encompassing the putative LFA-1 binding sites, fused to the Fc region of human IgG1 (*Pf*GBP-Fc) was generated (Fig. 3A). The *Pf*GBP-Fc fusion protein was engineered to ensure stability and solubility for binding assays, while the human IgG1 Fc tag facilitated detection via secondary anti-human IgG antibodies.

**Figure 3:**
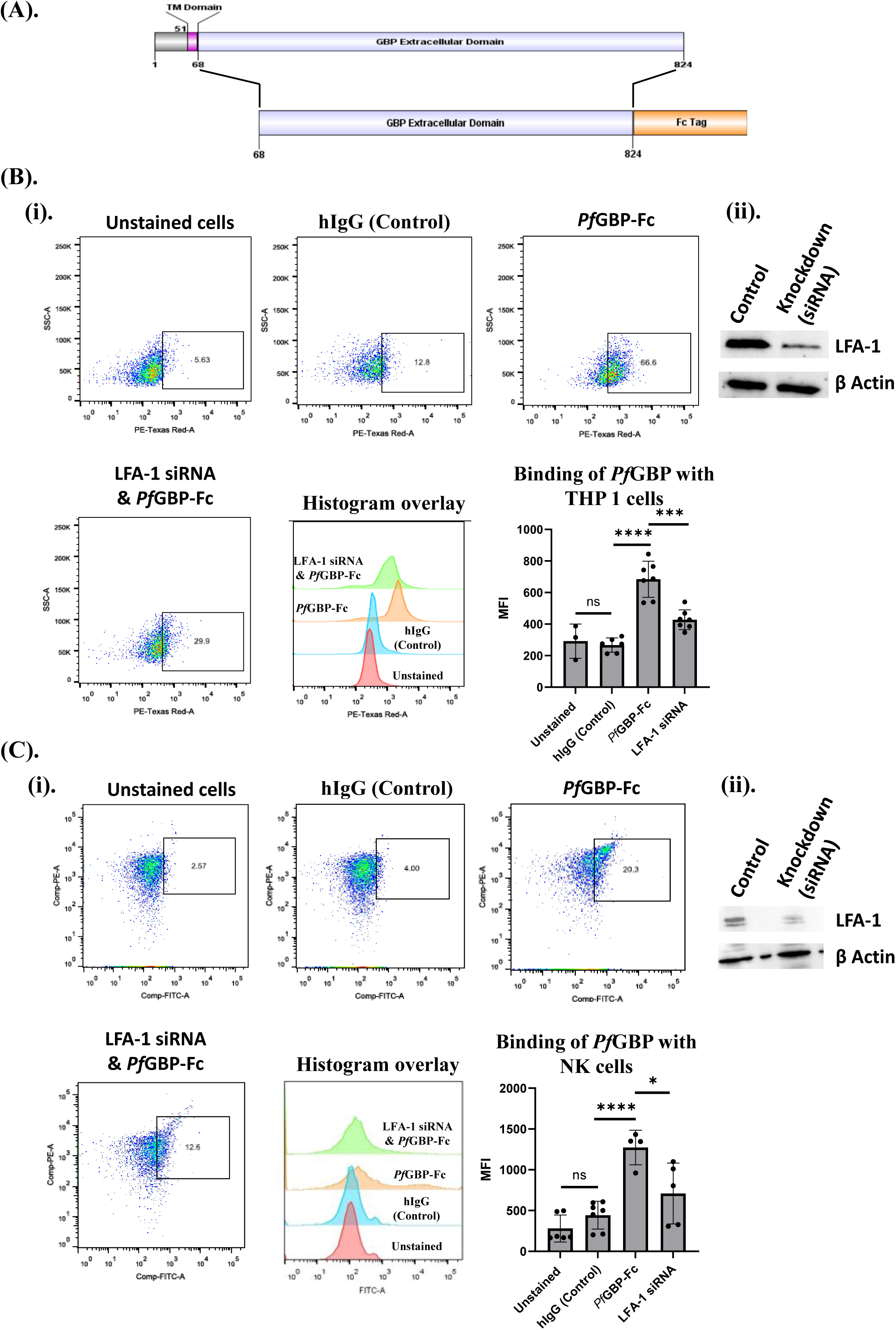
**Specificity of interaction between LFA-1 on Primary NK cells and THP-1 cells with *Pf*GBP-130**. Primary NK and THP-1 cells treated with LFA-1 siRNAs showed reduced binding to recombinant extracellular domain of *Pf*GBP-130 expressed in CHO K1 cells. **(A**) Schematics representation of *Pf*GBP-130 and its extracellular domain cloned in pFUSE-hIgG1-Fc2 vector for the expression in CHO K1 cells. The construct comprises the extracellular domain of *Pf*GBP-130 (including putative LFA-1 binding sites) fused to the Fc region of human IgG1. The Fc fusion provides stability, solubility, and facilitates detection. **(B)** Interaction of *Pf*GBP-130 ECD-Fc with THP-1 cells. **(i)** Flow cytometric analysis shows strong binding of *Pf*GBP-130 ECD-Fc to THP-1 cells, compared to an hIgG1 isotype control, indicating specific interaction. LFA-1 knockdown in THP-1 cells via siRNA treatment significantly reduced PfGBP-130 ECD-Fc binding, thus confirming the specificity of interaction between *Pf*GBP130 ECD-Fc and LFA-1 on THP cells. **(ii)** Western blot analysis using anti-CD11a antibody confirmed the siRNA-mediated knockdown of CD11a subunit of LFA-1 protein on THP-1 cells. **(C)** Interaction of *Pf*GBP-130 ECD-Fc with primary human NK cells. **(i)** Flow cytometry revealed a marked increase in *Pf*GBP-130 ECD-Fc binding to NK cells over the isotype control. siRNA-mediated knockdown of LFA-1 in NK cells led to a notable reduction in *Pf*GBP-130 ECD-Fc binding, confirming that LFA-1 is essential for this interaction. (**ii)** Western blot analysis using anti-CD11a antibody confirmed the siRNA-mediated knockdown of CD11a subunit of LFA-1 protein in NK cells, confirming LFA-1 as a critical receptor for *Pf*GBP130 ECD-Fc binding to both NK as well as THP-1 cells. * Denotes p < 0.05, ** denotes p < 0.01, and *** denotes p < 0.001. Representative flow plots depict the percentage of cells within a predefined positive gate, whereas the accompanying summary graph quantifies fluorescence intensity across the analyzed population. These two metrics report distinct properties of the distribution and are therefore not expected to be numerically identical.

The *Pf*GBP-Fc construct was transiently expressed in CHO K1 cells cultured in serum-free medium to minimize potential interference from serum components. The secreted fusion protein was subsequently purified from the culture supernatant using protein-A affinity chromatography (Suppl. Fig. 3A & 3B). The interaction between *Pf*GBP_-Fc and THP-1 was investigated, focusing on the role of LFA-1 as a potential receptor using siRNAs corresponding to CD11a subunit of LFA-1 mRNA. Next the cultured THP-1 monocytic cells with and without siRNA treatment were fixed and incubated with recombinant *Pf*GBP-Fc or control hIgG in FACS staining buffer for 2 hours. After washing, cells were stained with fluorescent anti-human PE-Texas Red antibodies (invitrogen) for binding analysis via flow cytometry to detect the binding of *Pf*GBP-Fc. Flow cytometric analysis demonstrated a significant increase (3-fold) in binding of *Pf*GBP -Fc compared to an hIgG isotype control to THP-1 cells, indicating a specific interaction of *Pf*GBP to THP-1 cells (Fig. 3B, i). To know whether the *Pf*GBP-Fc binding to THP-1 cells is LFA-1 dependent, we next performed Smartpool Accel siRNA-mediated gene silencing of CD11a subunit of LFA-1 in THP-1 cells (Suppl. Fig. 4B). As shown in figure 3B (ii), western blot analysis confirmed efficient depletion of the CD11a subunit of LFA-1 in THP-1 cells. Subsequent flow cytometric analysis revealed a substantial reduction in *Pf*GBP130-Fc binding to LFA-1-deficient THP-1 cells compared to untreated controls (Fig. 3B. i). This reduction thus definitively established LFA-1 as a crucial mediator of *Pf*GBP-Fc binding to THP-1 monocytic cells.

We further extended these findings to other immune cell populations, such as primary NK cells. Consistent with the results obtained in THP-1 cells, incubation of NK cells with *Pf*GBP-Fc resulted in a significant increase in binding intensity relative to the hIgG1 isotype control (Fig. 3C i). Smartpool Accel siRNA-mediated downregulation of CD11a subunit of LFA-1 in NK cell (Fig3C-ii & Suppl. Figure 4A), reduced *Pf*GBP130-Fc binding to NK cells (Fig. 3C. ii), thereby confirming the role of LFA-1 as a critical receptor for *Pf*GBP-Fc binding to NK cell.

LFA-1 predominantly expresses on immune cells, to further confirm ligand specificity of LFA-1, we evaluated the binding of *Pf*GBP130-Fc to multiple non-immune cell types, including HEK293T cells, HepG2 cells, and adipose derived stem cells (ADSC), which exhibit no or only very low basal expression of LFA-1. THP-1 cells, which express LFA-1, were included as a positive control. In contrast to THP-1 cells, all three non-immune cell types displayed minimal to negligible binding of *Pf*GBP130-Fc (Suppl. Figure 5A-D). Notably, HepG2 cells and stem cells showed only background-level binding comparable to the hIgG isotype control. These findings support that *Pf*GBP130-Fc binding is LFA-1 dependent and restricted to immune cells expressing LFA-1, reinforcing the specificity of the interaction.

### Engineered chimeric GBP expressing CHO cell line activates NK cell through LFA-1 binding

To elucidate the mechanism by which *P. falciparum* infected red blood cells (iRBCs) activate natural killer (NK) cells through LFA-1 engagement, we hypothesized that *Pf*GBP130 serves as a cognate ligand for LFA-1 on the iRBC surface. To test this hypothesis, a Chinese Hamster Ovary (CHO) cell line was engineered for the stable expression of *Pf*GBP-130 extracellular domain on its surface by leveraging lentiviral transduction. The engineered CHO-K1 expressed the extracellular C-terminal domain of *Pf*GBP-130 fused to the transmembrane domain of transferrin receptor (TfR-TM) for the surface presentation (Suppl. Fig. 3C). The surface expression was verified using anti-*Pf*GBP 130 antibodies by immunofluorescence (Fig. 4A. i). Flow cytometry analysis demonstrated robust binding of recombinant LFA-1 αI-Fc to *Pf*GBP-130–expressing CHO cells, while no significant binding was observed with mock-transduced CHO cells, confirming the specificity of the interaction (Fig. 4A. ii). To further investigate the role of *Pf*GBP-130 in NK cell activation and degranulation via direct engagement with its cognate receptor LFA-1, we co-cultured CHO cells stably expressing the extracellular domain of *Pf*GBP-130 (CHO–*Pf*GBP-ECD) with primary human NK cells at a 2:1 ratio (CHO–*Pf*GBP-ECD: NK). Given that LFA-1 affinity is dynamically regulated through inside-out signaling, co-cultures were stimulated with Poly I:C/ Lipofectamine 2000 (synthetic double-stranded RNA analogue), to induce a high-affinity conformational state of LFA-1. Subsequently, NK cell activation was evaluated by flow cytometric detection of early activation markers CD69 and CD25.Compared to mock CHO cells, *Pf*GBP130 expressing CHO cells elicited a significant ∼50% increase in CD69 and ∼40% increase in CD25 expression, indicating enhanced NK cell activation by *Pf*GBP-130. This activation was specific as addition of anti-CD11a antibodies in co-cultures reduced this activation significantly (Fig. 4B).

**Figure 4.**
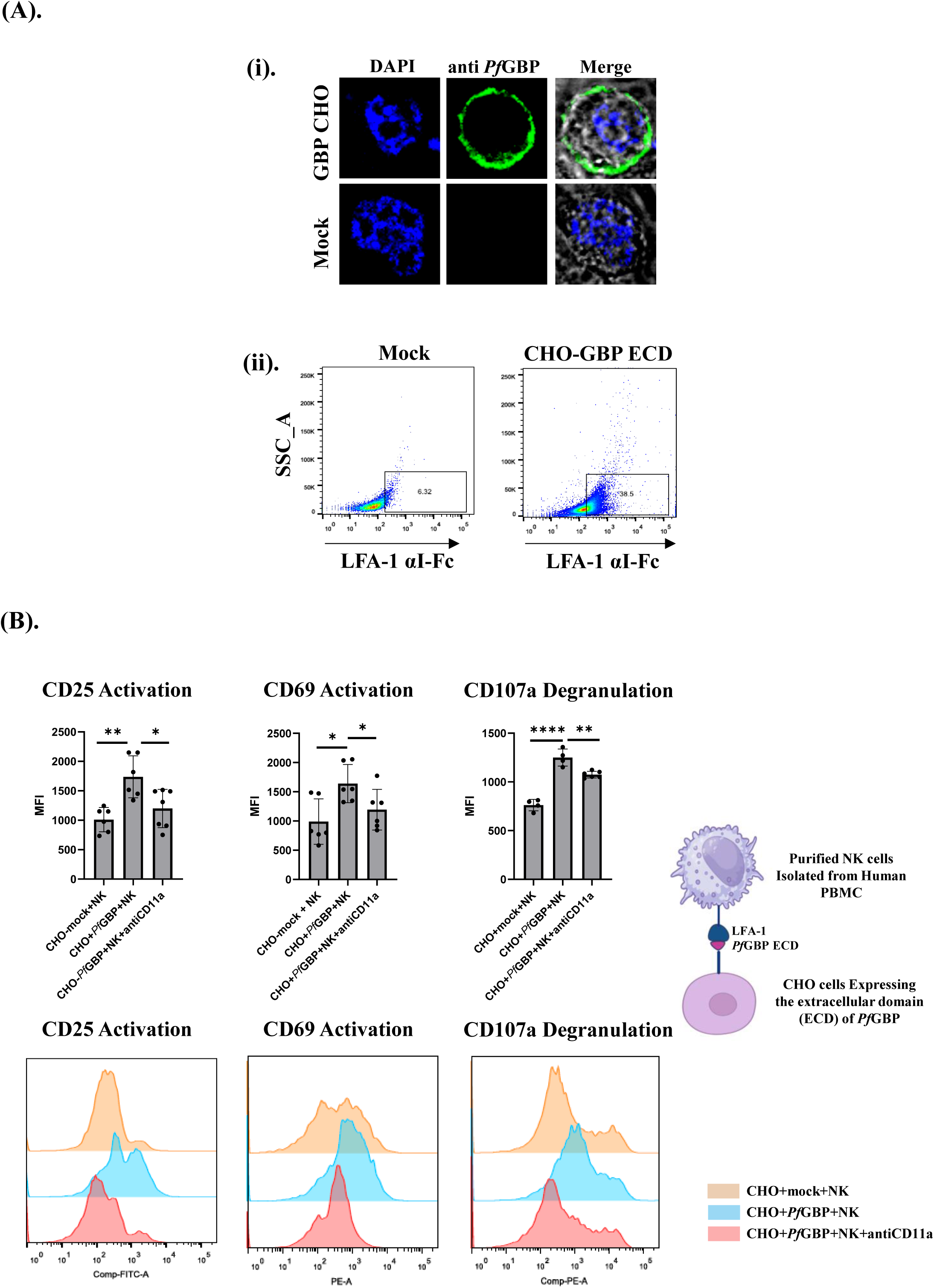
NK cells activated in the presence of *Pf*GBP-130. (A). **(i)** Expression of *Pf*GBP-130 ECD fused with Transferrin membrane domain on the membrane of CHO K1 cells by infecting the lentiviral vector; pMSCV Puro and its immunofluorescence analysis using anti-rabbit *Pf*GBP antibody. **(ii)** CHO K1 cells expressing *Pf*GBP-130 ECD bind LFA-1 αI-Fc. Binding of purified LFA-1 αI-Fc to *Pf*GBP-130 ECD expressing CHO cells was assessed by FACS using an PE-Texas red anti-human IgG antibody. **(B)** NK cells activation in the presence of CHO K1 cells expressing *Pf*GBP ECD. Human NK cells were purified (>95%) from fresh PBMC and co-cultured with CHO K1 cells expressing *Pf*GBP ECD in 2: 1 ratio (20,000 CHO-K1 cells:10,000 NK cells) and these cells were stimulated with (Poly I:C/lipofectamine 2000) for 24 h. NK cells were separated from adherent CHO K1 cells and NK cells activation was assessed by assaying the expression of activation markers (CD69, CD25) and a degranulation marker; CD107a . NK cells co-cultured with CHO K1 cells expressing *Pf*GBP-ECD protein showed significant increase in the expression of CD25 and CD69, as well as CD107a in comparison to the NK cells co-cultured with mock CHO cells. Addition of anti-CD11a (HI111 clone) antibodies reduced the expression of both activation and degranulation markers. * Denotes p < 0.05, ** denotes p < 0.01, and *** denotes p < 0.001.

Furthermore, we examined the impact of *Pf*GBP-130 expression on primary NK cells degranulation, an indirect indicator of cytotoxicity. For this surface expression of CD107a, a marker of lysosomal degranulation was measured using flow cytometry. Co-culture of NK cell with CHO–*Pf*GBP-ECD resulted in a substantial increase in CD107a on primary NK cells as compared to mock CHO control. This increase was significantly reduced in the presence of anti-CD11a antibodies, indicating that the enhanced degranulation was specifically mediated through LFA-1 engagement with *Pf*GBP-130 on the CHO cell surface (Fig. 4B). Together, these results indicate that *Pf*GBP-130 protein promotes LFA-1 mediated activation and cytotoxic degranulation of NK cells.

### Human NK cells eliminate iRBCs in a co-culture study and this elimination is dependent on GBP-130 expression on iRBCs

To gain direct evidence of NK cell mediated killing of iRBCs via *Pf*GBP-130 dependent interaction, we performed co-culture experiments using purified human NK cells and iRBCs as described earlier by Chen at al 2014 (Chen et al., 2014), in the presence or absence of anti-*Pf*GBP-130 antibodies. An isotype control antibody served as a negative control for non-specific antibody effects. Briefly, isolated human NK cell was treated with Fc receptor blocker using anti-CD16 (3G8 clone) prior to addition of iRBC (0.5% parasitemia) in 10: 1 ratio (NK: iRBC). The blocking of CD16 with anti CD16 (clone 3G8) inhibits Fc engagement with CD16 receptor and thereby abrogates any potential antibody dependent cellular cytotoxicity (ADCC) contribution during the NK-iRBC co-culture in presence of anti GBP neutralizing antibody (Mandelboim et al., 1999; Yeap et al., 2016). The co-cultures were maintained for 48 hour and 96 hours, a time frame sufficient for NK cell activation and parasitemia control respectively.

In a parallel set of experiments, we sought to specifically examine the effect of blocking *Pf*GBP-130 on iRBC with anti-*Pf*GBP neutralizing antibody on NK cell activation and parasitemia control. The purity of isolated NK cells was accessed using anti-CD3 and anti-CD56 (Suppl. Figure 3D). Isolated NK cells were co-cultured with iRBCs for an extended period of 96 hours to allow for a more comprehensive assessment of parasite growth, which was assayed with Hoechst staining and number of NK cells were assessed by anti-CD56 NK antibody staining in a flow cytometry analysis. The iRBCs were positive for Hoechst, but negative for CD56, while NK cells were positive for both. The co-culture of NK cells with iRBC either alone or in presence of rabbit IgG (rIgG) isotype control antibody, significantly reduced parasitemia compared to iRBCs cultured in the absence of NK cells. In contrast, the addition of anti-*Pf*GBP antibody to NK cell-iRBC co-cultures resulted in a marked and statistically significant increase in parasitemia, indicating that blockade of *Pf*GBP-130 impairs NK cell–mediated parasite clearance (Fig. 5A). Conversely, parasitemia levels in the co-cultures supplemented with the isotype control antibody were observed to be comparable to those in the NK cell culture alone, indicating that the observed reduction in parasitemia control was specifically attributable to the blockade of *Pf*GBP-130 and not due to non-specific antibody interactions (Fig. 5A). This finding confirms the specificity of the anti-*Pf*GBP antibody and reinforces the critical role of *Pf*GBP in NK cell-mediated parasite clearance.

**Figure 5.**
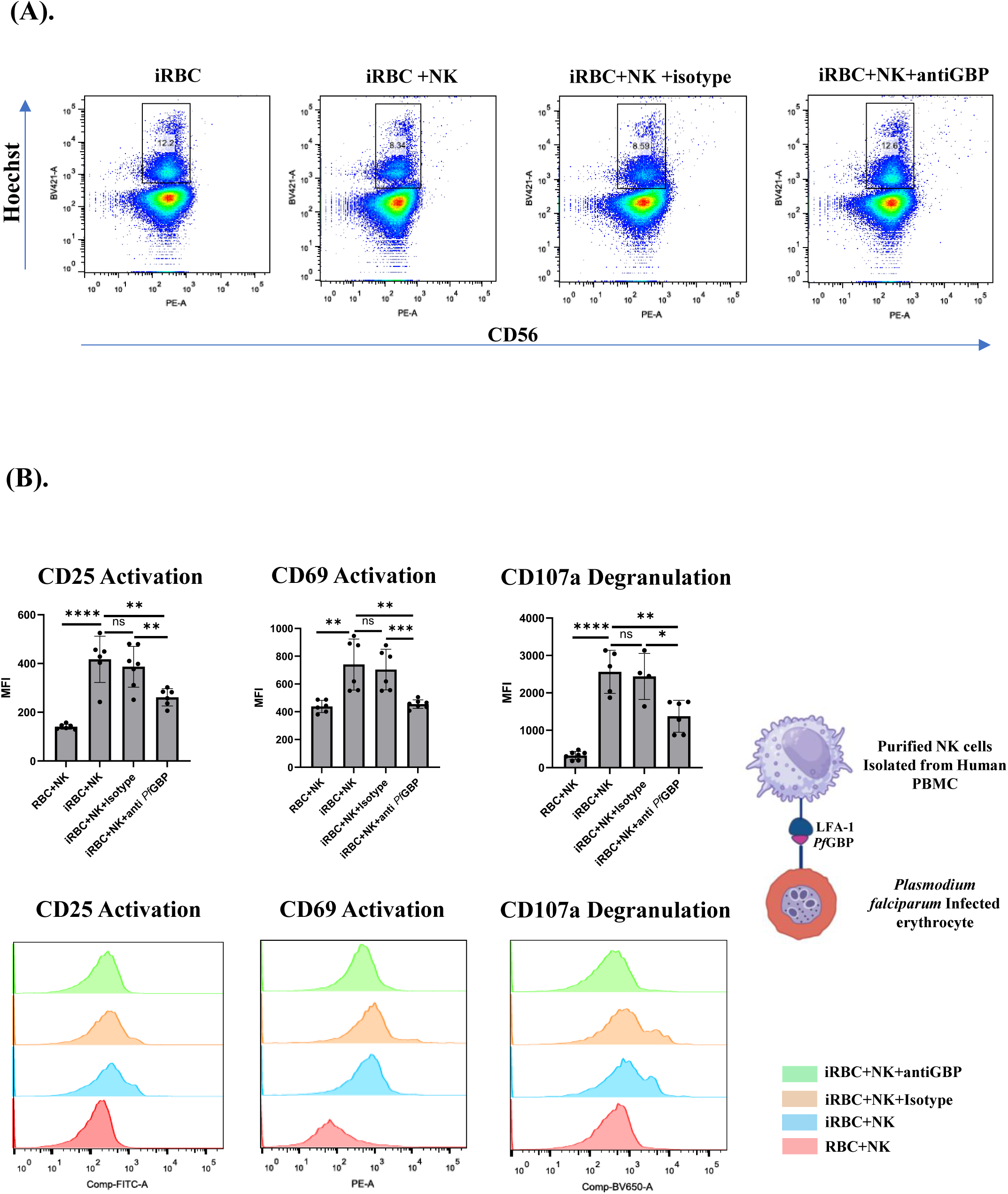
NK cells activated in the presence of iRBCs, controls parasite infection **(A)** Activated human NK cells eliminate iRBCs *in vitro.* Human NK cells when co-cultured with iRBCs reduce parasitemia significantly after 96h and in the presence of anti-*Pf*GBP-130 antibodies this reduction in parasitemia was blocked. Presence of anti-GBP-130 abs resulted in parasitemia simialr to the control when NK cells were incubated with iRBCs alone. **(B)** NK cells activation in the presence of iRBCs. Human NK cells were were purified (>95%) from fresh PBMC and co-cultured with synchronized schizont stage iRBCs at a parasitemia of 0.5% in 10:1 ratio (NK: iRBC) for 48h. Quantification of activation and degranulation markers was performed after 48 hours. NK cells co-cultured with iRBCs showed significant increase in the expression of CD25 and CD69, the two activation markers as well as for the expression of CD107a, a degranulation marker in comparison to the NK cells co-cultured with RBCs alone. Addition of anti-rabbit PfGBP-130 antibodies reduced the expression of both activation and degranulation markers in these NK cells in comparison to rabbit IgG isotype control. * Denotes p < 0.05, ** denotes p < 0.01, and *** denotes p < 0.001.

We further studied the levels of activation and degranulation markers in NK cell in the absence and presence of anti-GBP-130 antibody (Fig. 5B). Briefly, NK cell and iRBC were co-cultured in the presence to anti-GBP-130 antibody or rabbit IgG (rIgG) isotype control antibody. Incubation of human NK cells with iRBC for 48 hrs showed a significant percent change in MFI of activation markers; CD69 and CD25 as well as degranulation marker; CD107a. This effect was reduced in the presence of anti-*Pf*GBP antibody, which signified that *Pf*GBP-130 engagement is necessary for the activation and degranulation of NK cell (Fig. 5B). These results were specific as levels of activation and degranulation markers were same in NK cells incubated with iRBC in the presence or absence of rIgG control. Taken together, the reduced NK cell activation and increase in parasitemia following *Pf*GBP-130 blockade strongly support the conclusion that direct interaction between LFA-1 on NK cells and GBP-130 on iRBCs is a critical determinant of contact dependent NK cell-mediated immune responses. These findings further establish *Pf*GBP-130 as a key ligand in the LFA-1–dependent activation of NK cell during *Plasmodium falciparum* infection.

## Discussion

NK cells have been well described for their ability to eliminate viral-infected cells, tumors and several pathogens including *P. falciparum* (Burrack et al., 2019). A fine-tuned panel of activating and inhibitory receptors regulates the activation of NK cells. Among these, LFA-1, an integrin family member, serves not only as an adhesion molecule but also as an important activating receptor on NK cells (Urlaub et al., 2017). Beyond mediating adhesion, LFA-1 signaling is essential for the polarization of lytic granules towards target cells, a prerequisite for efficient NK cell cytotoxicity(Bryceson et al., 2005; Kabanova et al., 2018).

There are mounting evidence to show that NK cells contribute to immune responses in the clearance of parasites, removal of infected hepatocytes and infected RBCs through cytotoxicity and ADCC (Burrack et al., 2019). Using an immunodeficient humanized RAG-IL2Rγc–deficient (RICH) mice, a study by Chen et. al., showed that NK cells, rather than macrophages are responsible for regulating the parasite growth *in vivo* by directly interacting with iRBCs (Chen et al., 2014). This study also implicated LFA-1 as key receptor involved in NK cell mediated killing of iRBCs. LFA-1 is a heterodimer composed of CD11a/CD18, also called α1/β2. LFA-1α subunit,αL has two prominent structural features; I domain of 200 aa residues and three EF hand-like domains, which are crucial for ligand binding. Two other integrins expressed on leukocytes; Mac-1 (CD11b/CD18) and p150.95 (CD11c/CD18) share the same CD18 or ß2 integrin subunit and have homologues α subunits (Kolanus et al., 1996). The I domain of these proteins are homologous to motifs in von Wliilibrand factor, cartilage matrix protein, collagen type VI, complement factor C2 and factor B (Huang and Springer, 1995). Structure-function studies on LFA-1 and Mac-1 have implicated their I domains in ligand binding (McDowall et al., 1998). Here, we set out experiments to know whether LFA-1 αI, a 200 aa domain of LFA-1 on NK cells alone recognizes the iRBCs. To do so, LFA-1α1 domain was expressed as a fusion protein with Fc domain of hIgG and was allowed to bind *P. falciparum* infected erythrocyte. LFA-1α1-Fc fusion protein bound significantly with iRBCs in comparison with hIgG control protein.

Having established LFA-1α1 as a key binding domain to iRBCs, we next carried out studies to identify the parasite derived ligand on iRBC surface that interacts with LFA-1. Up till now, LFA-1 has been shown to bind human and mouse ICAM-1 (Huang and Springer, 1995). A unique property of LFA-1 is that its binding affinity can be dynamically modulated by inside-out (intracellular signals triggered by antigens and chemokines) and outside-in (ligand binding) signaling pathways (Gerard et al., 2021; Kondo et al., 2022). Conformation changes and membrane clustering of LFA-1 have been shown to significantly influence its binding affinity (Shi and Shao, 2023; Urlaub et al., 2017). To fish out the parasite ligand on the surface of infected RBCs for LFA-1, here we cross-linked LFA-1α1-Fc fusion protein with iRBCs, followed by immunoprecipitation and LC-MS/MS analysis. *Pf*GBP-130 protein was identified as a prominent LFA-1interacting protein on the iRBC surface. As the name suggests, *Pf*GBP-130 is a glycophorin binding protein consisting of 11 highly conserved 50 amino acid repeats and a charged N-terminal region of 225 amino acids (Kochan et al., 1986). A number of studies have shown that *Pf*GBP-130 is exported out of iRBC surface and is involved in erythrocyte invasion (Soni et al., 2016). We further confirmed the iRBCs surface expression of *Pf*GBP-130 protein using anti-*Pf*GBP antibodies, thus supporting the observation that it probably is involved in recognition of iRBCs by NK cells. Western blot analysis of LFA-1 αI -Fc bound iRBCs lysate using anti-*Pf*GBP antibodies further strengthened the hypothesis that *Pf*GBP-130 on iRBCs surface is the ligand for LFA-1 that helps NK cells to recognize the iRBCs. To evaluate the specificity of LFA-1 interaction with *Pf*GBP-130 on iRBC surface, we overexpressed external domain of *Pf*GBP130 as a Fc fusion protein and evaluated its binding to THP-1 monocytes and primary NK cells. Recombinant *Pf*GBP-Fc protein efficiently bound to the surface of both cell types. Importantly, siRNA-mediated knockdown of CD11a subunit LFA-1 significantly reduced this binding, confirming that *Pf*GBP-130 directly interacts with LFA-1 on immune cells, thereby advocating GBP-130 as a ligand for LFA-1.

NK cells are predominantly cytolytic lymphocytes, and their primary effector function relies on the degranulation of cytotoxic molecules. LFA-1 not only mediates tight adhesion to target cells but also plays a crucial role in early NK cell activation and degranulation (Barber et al., 2004). To assess whether *Pf*GBP-130 triggers NK cell activation through LFA-1 engagement, we analyzed the expression of activation markers CD25 and CD69, and the degranulation marker CD107a, in NK cells co-cultured with CHO cells expressing the extracellular domain of *Pf*GBP-130. Compared to mock controls, co-culture with *Pf*GBP-130–expressing CHO cells significantly upregulated all three markers. Importantly, this activation was abrogated by the addition of anti-CD11a antibodies, confirming that *Pf*GBP-130 mediates NK cell activation and degranulation via LFA-1. CD107a (LAMP-1), a lysosomal membrane protein, serves as a well-established marker of NK cell degranulation (Urlaub et al., 2017; Ye et al., 2018).

Besides mediating the tight adhesion of NK cells to various target cells, LFA-1–mediated signaling plays a pivotal role in the polarization of lytic granules towards the immunological synapse, thereby enhancing NK cell–mediated cytotoxicity (Barber et al., 2004; Shi and Shao, 2023). To know whether such a toxicity is also generated as a result of interaction between NK cells and iRBCs, we analyzed the expression of two activation markers; CD25 and CD 69 and a degranulation marker; CD107a in NK cells co-cultured with CHO cells overexpressing *Pf*GBP-130 protein on their surface as described by Ye, W et al. 2018 (Ye et al., 2018). All these three markers showed significant upregulation in primed NK cells in comparison to naïve NK cells. This upregulation was specific as the presence of anti-LFA-1 antibodies reversed these upregulations. To determine whether the activation of primary NK cells is specifically mediated by direct engagement between LFA-1 on NK cells and *Pf*GBP-130 on iRBCs at the immunological synapse, we performed targeted co-culture assays. we co-cultured purified human NK cells with iRBCs and measured both parasite growth as well as the levels of activation and degranulation markers in NK cells, in the presence or absences of anti-*Pf*GBP-130 antibodies. The results of co-cultured study showed that the presence of iRBCs resulted in activation of NK cells and this resulted in reduced parasitemia. This activation of NK cells was as a result of *Pf*GBP-130 expression on iRBCs as anti-*Pf*GBP-130 antibodies reduced the activation of NK cells. These results are in line with a recent study that showed upregulation of these markers in NK cells co-cultured with iRBCs from human population (Arora et al., 2018; Ye et al., 2018).

In conclusion, the results presented here identify *Pf*GBP-130 as a novel ligand for LFA-1, in particularly its LFA-1 αI domain on NK cells. This interaction mediates firm adhesion between human NK cells and *Plasmodium falciparum* infected red blood cells resulting in the activation of NK cells. The ensuing activation subsequently triggers NK cells cytotoxic degranulation resulting in targeted killing of iRBCs. Given that NK cells constitute first line of defense against malaria, this molecular mechanism appears to be important in early control of *P. falciparum* infection in particularly by implementing a host directed therapy.

## Materials and Method

### *Plasmodium* culture of 3D7

*Plasmodium falciparum* 3D7 parasites were maintained in O+ human erythrocytes at 4% hematocrit in RPMI 1640 medium (pH 7.4) supplemented with 50 mg/l hypoxanthine, 5% Albumax II, 2 g/l sodium bicarbonate, and 20 µg/ml gentamycin according to the protocol described by Trager and Jensen (Trager and Jensen, 1976). Cultures were maintained at 37 °C in a gas mixture of 90% N2, 5% CO2, and 5% O2. Parasite synchronization was performed using 5% (w/v) sorbitol.

### Cell lines and Primary Human NK cell culture

THP-1 cells, CHO-K1, and HEK293T cells were cultured in RPMI 1640 (Invitrogen) supplemented with 10% fetal bovine serum (FBS), or in DMEM (for HEK293T) with 10% FBS. Primary human NK cells were isolated from peripheral blood mononuclear cells (PBMCs) by negative selection using the BioLegend NK Cell Isolation Kit, achieving >95% purity. Adipose derived stem cell (ADSC) was cultured in commercial serum free media MesenCult™-XF as manufacturer instruction.

### Fusion protein construct and its expression and purification

The LFA-1 αI domain was PCR-amplified from THP-1 cDNA and cloned into the pFUSE-hIgG Fc2 vector (InvivoGen). The construct was transfected into CHO-K1 cells using JetPrime (Polyplus). Post-transfection, cells were maintained in IgG-depleted FBS-containing media and later switched to 10% Nuserum medium. After 72 h, the culture supernatant was collected, filtered, and the secreted fusion protein purified using Protein A agarose affinity chromatography (G-Biosciences). Proteins were concentrated using 3 kDa Centricon filters (Merck) and quantified via BCA assay.

### Immunoprecipitation and Mass spectrometry

Schizont-stage *P. falciparum* iRBCs were isolated via Percoll gradient centrifugation and incubated with 100 µg LFA-1 αI-Fc or human IgG control for 2 h at 37 °C. Crosslinking was performed using 5 mM DTSSP (Thermo Fisher) for 30 min. Cells were lysed with IP Native Lysis Buffer and incubated with Protein A magnetic beads (Thermo Fisher) for 2 h. Beads were washed and proteins eluted with 5 mM DTT. Eluates were processed for LC-MS/MS analysis. Eluates were processed for LC-MS/MS analysis. The eluates were first precipitated using 10% trichloroacetic acid (TCA), 5% acetone, and 5 mM DTT. Samples were air-dried and resuspended in 100 mM ammonium bicarbonate (ABC) buffer containing 8 M urea, then diluted to 1 M urea with 100 mM ABC buffer. Followed by reduction with 10 mM DTT for 1 h at room temperature (RT), and alkylation with 40 mM iodoacetamide (Sigma-Aldrich) for 1 h at RT in the dark. The alkylated eluates were further digested with trypsin (1:50, wt/wt) at 37 °C for 12–16 h. Peptides were acidified with 0.1% formic acid and analyzed on an Orbitrap Fusion Lumos mass spectrometer coupled to a Nano-LC 1200 (Thermo Fisher Scientific). Data were processed with Proteome Discoverer v2.4 using the SEQUEST and AMANDA algorithms.

### iRBC and Immune cell binding assay

*P. falciparum* 3D7 parasites were cultured to 5% parasitemia and synchronized. Ring, trophozoite, and schizont stages were isolated using Percoll gradient. Approximately 10[iRBCs from each stage were incubated with 250 nM purified LFA-1 αI-Fc fusion protein or 250 nM human IgG (hIgG) control for 2 h. After washing, cells were stained with PE-Texas Red or FITC-conjugated anti-human Fc antibodies (Invitrogen) for 30 min and analyzed by flow cytometry. Similarly, THP-1 cells and primary human NK cells, including siRNA-treated populations, were fixed and incubated with GBP130-Fc (250 nM) or hIgG control (250 nM) in FACS buffer for 2 h. After washing, cells were stained with PE-Texas Red–conjugated anti-human secondary antibodies and analysed by flow cytometry.

### Cloning, expression, and purification of *Pf*GBP130 (PF3D7_1016300)

PF3D7_1016300 (aa 69–270; encompassing one GBP repeat) was amplified from *P. falciparum* genomic DNA/cDNA, cloned into pGEM-T, and confirmed by restriction digestion and sequencing (Suppl. Figure 2D). The verified fragment was subcloned into pET28b between the NcoI/XhoI sites and transformed into *E. coli* BL21(DE3). Recombinant expression was induced with 1 mM IPTG for 5 h at 37°C, and cells were lysed by sonication in Tris-buffer (50 mM Tris, pH 8.0; 150 mM NaCl). Soluble and insoluble fractions were separated by centrifugation, and expression was assessed by SDS-PAGE and anti-His immunoblotting. His-tagged *Pf*GBP-130 was purified from lysates using Ni-NTA affinity chromatography, with wash steps in lysis buffer containing 10 mM imidazole and elution using a 50-500 mM imidazole gradient. Polyclonal antisera against purified recombinant fragments were raised in BALB/c mice and New Zealand White rabbits as described previously (Sachdeva et al., 2006).

### *In vitro Pf*GBP-130/LFA-1 **α**I-Fc interactions analysis with BLI, *in silico* docking and Molecular Dynamic stimulation analysis

The real-time interactions between LFA-1 and its potential binding partner, were studied using Bio-layer interferometry (BLI) on the Sartorius Octet K2 system. In this setup, LFA1 was immobilized onto an AR2G sensor (Octate Amine Reactive Second-Generation biosensor) at a concentration of 100ng, and the unbound protein was washed off using I X Phosphate-buffered Saline (PBS) pH7.4 buffer. To determine the binding affinity, *Pf*GBP was tested at various concentrations. Throughout the experiment, IX PBS pH7.4 was used as both the running and the dissociation buffer, and all working dilutions of *Pf*GBP proteins were prepared in the same buffer. For data normalization, a control sensor with immobilized LFA1 was run in parallel, where only the running buffer (1X PBS) flowed over the sensor. The interaction kinetics, including the association and dissociation curves, were monitored for 800 seconds in a series of increasing concentrations of the *Pf*GBP protein. The data was acquired with Octet BLI discovery 12.2 software, and the KD value was calculated through the Octet Evaluation software, applying a 1:1 binding model for fitting the curve. The experiment was performed at 25°C.

*In silico* docking and simulation, approaches were employed to investigate the molecular interaction between LFA1 with *Pf*GBP. To achieve this, initially, a homology model for PfGBP was generated using Alphafold(Jumper et al., 2021), and the quality of the generated model was assessed with PROCHECK and PyMOL. While for LFA1, its crystal structure (PDB code: 1LFA) was employed in the interaction studies. Next, rigid body protein-protein docking was carried out using the Cluspro2.0 web server (Comeau et al., 2004), a top performer at CAPRI (Critical Assessment of Predicted Interactions) challenges. ClusPro ranked the docked models based on cluster size and energy. The default set of docking parameters was applied, and the top-ranked complexes were selected based on the largest cluster sizes and lowest energy values. The number of cluster member and the model cluster score (that is cluster center and lowest energy) are shown in Suppl Table 2. The cluster center weighted score shows the structure in the cluster that has the highest number of neighbour structures, whereas the lowest energy score represents the structure which has the lowest energy in the cluster.

The models were further analyzed to examine the interaction interface using PyMOL (PyMOL Molecular Graphics System, v2.1 by Schrödinger, LLC). The final docked model for the LFA1/*Pf*GBP complex was chosen based on significant interactions and the model with the heights buried surface area (BSA). PDBsum software was utilized to analyze the interactions within the complex(Laskowski et al., 2018).

To examine the stability, the LFA1/*Pf*GBP complex was subjected to molecular dynamics (MD) simulations using the Desmond module of SchroCdinger software. Firstly, the Schrödinger protein preparation tool was used to prepare the protein; by removing the non-bonded water (> 3Å from protein residues), optimizing the H-bonds, and energy minimizing the final structures using OPLS3. Next, these complexes were placed in an orthorhombic water box at a buffer distance of 10 Å, and solvated using the TIP4PEW solvation model while maintaining a NaCl concentration of 0.15 M to ensure a physiological ionic strength. After solvation, the complex was subjected to energy minimization using OPLS4 force field parameters followed by relaxation, and the simulation was performed for 200 ns at NPT conditions with a frame captured at every 0.2 ns. For determining the trajectories, the frames were collected and examined through a simulation interaction diagram, which helped in determining fluctuations (RMSD). The final frame of the simulation was saved in a PDB file format. Further, PDBePISA (Protein Interfaces, Surfaces, and Assemblies (PISA) server was employed to analyze the intermolecular interfaces(Krissinel and Henrick, 2007), including computing buried surface area (BSA Å) for the complex. PyMol was used to represent these interactions, while binding energy estimations were performed employing multiple computational tools (Sukhwal and Sowdhamini, 2015; Weng et al., 2019; Xue et al., 2016; Yang et al., 2023).

### Chimeric CHO stable cell line generation

To generate CHO-K1 cells stably expressing *Pf*GBP130, the codon-optimized synthetic gene encoding the extracellular domain of *Pf*GBP130 was cloned into the lentiviral transfer plasmid pMSCV-puro. Lentiviral particles were produced by co-transfecting HEK293T cells with pMSCV-puro-*Pf*GBP130, psPAX2 (packaging plasmid), and pMD2.G (VSV-G envelope plasmid) using JetPrime transfection reagent (Polyplus), following the manufacturer’s instructions. After 72 h, viral supernatants were harvested and concentrated using Lentivirus Concentrator Solution (Takara). CHO-K1 cells were transduced with the concentrated lentivirus for 48 h to facilitate viral entry and integration of *Pf*GBP 130 gene, and selected with puromycin (5 µg/mL) for 7 days to enrich for stably transduced cells. Following antibiotic selection, cells were expanded in fresh culture medium, and *Pf*GBP130 surface expression was confirmed by fluorescence microscopy.

### NK cell activation assay

*Pf*GBP130-expressing CHO stable cells were trypsinized and counted before being co-incubated with purified primary human NK cells at a 2:1 ratio (CHO: NK). Following this co-incubation, a Lipofectamine/PolyI:C complex (100 µg/mL) was added to each well to stimulate a response for 24hrs. To investigate the role of LFA-1, NK cells were pre-incubated with a neutralizing antibody (anti CD11a, clone HI111) against the LFA-1 receptor for 30 minutes at room temperature before being added to the *Pf*GBP130-CHO cells.

### siRNA mediated knockdown of THP1 and NK cells

siRNA mediated CD11a knockdown was performed in both THP-1 cells and primary human NK cells using Accell SmartPool siRNA (Dharmacon). THP-1 cells were seeded at 1[×[10[cells/well in 96-well plates in Accell siRNA transfection medium containing 2.5% FBS and 1µM CD11a-targeting siRNA. Primary NK cells (1[×[10[) were suspended in Accell siRNA transfection medium supplemented with 10 ng/mL IL-15 and transfected with 2 µM CD11a siRNA. Cells were incubated for 72 h, after which knockdown efficiency was confirmed by immunoblotting with anti-CD11a antibodies.

### Flowcytometry

The following antibodies were used for staining: anti-human CD3 (biotin, OKT3; Elabsciences), anti CD16 (3G8; Biolegend), anti CD11a (HI111; BioLegend), CD56-PE (5.1H11; Elabsciences), Ultra-LEAF™ Purified Human IgG1 Isotype (Biolegend), Goat anti-Human IgG Alexa Fluor™ 488 (Invitrogen), Goat anti-Human IgG Secondary Antibody, Alexa Fluor™ 594 (Invitrogen), (CD107a (H4A3; Elabsciences), CD25-Alexa Fluor® 488 (BioLegend), and CD69 (FN50; Elabsciences). Purified NK cells, either alone or co-cultured with iRBCs, were stained in 100 µL PBS containing 0.2% BSA and 0.05% sodium azide for 30 min on ice. After washing, stained cells were analyzed on a BD LSR Fortessa X-20 flow cytometer, and data were processed using FlowJo software (BD Biosciences).

### NK cell co-culture with RBC and iRBC

Human NK cells were purified from PBMCs using the BioLegend Human NK Cell Isolation Kit, achieving >95% purity. Prior to co-culture with RBCs/iRBCs, NK cells were pre-incubated with Fcγ receptor blocker (anti-human CD16, clone 3G8; 2 µg per 10[cells) for 30 min at room temperature. Late-stage *P. falciparum* schizont-infected RBCs (iRBCs) were purified at 0.5% parasitemia. For neutralization experiments, either anti-*Pf*GBP130 antibody or isotype control IgG1 was added to the culture wells for 30 min prior to NK cell addition. Pre-treated NK cells were then co-cultured with iRBCs or uninfected RBCs at a 10:1 ratio (NK: iRBC or NK: RBC) for 48 h. For activation marker analysis, cells were collected and stained with relevant antibodies. For parasitemia assessment, co-cultures were maintained for 96[h. Cells were stained with Hoechst 33342 (20 µg/mL) and anti-human CD56-PE (BioLegend) for 15 min, washed, and analyzed using a BD LSR Fortessa X-20 flow cytometer.

### Statistical Analysis

Data are presented as mean ± SEM. Differences between groups were analyzed via Student *t* test. A P value <0.05 was considered statistically significant. All calculations were performed using GraphPad Prism 10 software package.

## Ethical Approval

The study was conducted according to the guidelines of the Declaration of Helsinki and approved by the International Centre for Genetic Engineering and Biotechnology (ICGEB)’s Scientific Ethical Review Unit and the Institutional Animal Ethics Committee.

## Funding and acknowledgements

The research work in PMs and AMs is supported by the Flagship Project (BT/IC-06/003/91), DBT-Wellcome; Team Science Grant (WTA/24/006); and JC Bose Grant (DST/20/015) from the Department of Science and Technology, Govt. of India. The funders had no role in the design of the study; the collection, analysis, and interpretation of the data; or the writing of the manuscript. We also thank the Rotary Blood Bank (India) for providing human red blood cells. We also acknowledge the Immunobiology group at ICGEB for providing infrastructure and insightful suggestion for the completion of this study.

## Author Contributions

O.M., R.D., performed the research experiments. S.S.S., contributed expertise and necessary tools for LC–MS studies. G.P., and A.P., performed the protein purification, antibody generation and immunolocalization assays related to *Pf*GBP. P.K., performed the BLI related experiments. N.P., M.A., M.M.I., A.T. and P.A., maintained *Plasmodium* Culture and provided assistance in immunolocalization studies. P.M., D.K., and A.M., designed the study, interpreted the data and wrote the manuscript; P.M., A.M., D.K., O.M., R.D., and A.P., contributed to the experimental design, data interpretation, and manuscript preparation.

## Supplementary Figures Legends

**Suppl. Figure 1.** Amino acid sequence alignments of “αI domain” of Lymphocyte Function Associated Antigen-1 (LFA-1) with similar Von Willebrand Factor (vWA) domains of human and Mac-1 I-domain showing the similarity and differences among these proteins.

**Suppl.** Figure 2**. (A)** Schematic showing Vector map of pFUSE-hIgG1-Fc2 and sites used for cloning the LFA αI domain gene used for the expression of LFA αI-Fc fusion protein. **(B)** Agarose gel electrophoresis showing PCR amplification of DNA corresponding to LFA αI domain and restriction analysis of cloned plasmid; pFUSE-hIgG1- LFA αI -Fc2. **(C).** Specificity of LFA-1 αI-Fc binding to infected erythrocytes. Importantly, no significant difference in binding to uRBCs was detected between LFA-1 αI–Fc and the isotype control, indicating minimal non-specific interaction with the erythrocyte surface. **(D) (i)** Agarose gels showing the PCR amplification of 600 bp fragment of *PfGBP-130* gene (Pf3D7_1016300). **ii. & iii.** Agarose gel electrophoresis showing restriction analysis of cloned *PfGBP-130* gene fragment in pGEM-T and pET-28b vectors respectively.

**Suppl.** Figure 3**. (A)** SDS-PAGE and **(B)** western blot analysis showing the purified *Pf*GBP130-Fc fusion protein expressed in CHO K1 line. Western blot analysis was performed using anti-IgG abs. **(C)** Schematics of cloned *Pf*GBP130-ECD fused to TfR-TM domain in pMSV puro plasmid to produce lentiviral vector. **(D)** Staining of purified primary NK cells using anti-CD3 and anti-CD56 showing purity of NK cells.

**Suppl.** Figure 4**. (A)** FACS analysis to show expression of LFA-1 subunit CD11a using anti CD11a antibody on primary NK cells and its expression inhibition on the same cells using Smart pool Accel siRNA corresponding to CD11a of LFA-1. **(B)** FACS analysis to show expression of CD11a subunit of LFA-1 on THP-1 cells and its expression inhibition on the same cells using Smartpool Accel siRNA corresponding to CD11a of LFA-1. Secondary anti-mouse PE-Texas Red was used for detection.

**Suppl.** Figure 5 **(A-D).** To confirm ligand specificity and exclude non-specific binding, *Pf*GBP130-Fc binding was assessed on multiple cell types by flow cytometry. THP-1 cells, which express LFA-1, were used as a positive control, while non-immune cell lines (HEK293T, HepG2) and stem cells (low/negative basal LFA-1 expression) were included as negative controls. Robust *Pf*GBP130-Fc binding was observed on THP-1 cells, whereas HEK293T, HepG2, and stem cells showed minimal to negligible binding comparable to the hIgG isotype control.

## Supplementary Tables

**Suppl. Table 1:** Table of MS/MS hits of the beads+hIgG control.

**Suppl. Table2**: Docking scores and binding energy calculated using various computational tools to assess the quality of the GBP-LFA1 docked complex.

