## Supplemental figures 1-5, and Supplemental table 1-2 for "LFA-1 Interaction with GBP-130 on *Plasmodium falciparum*-infected Red Blood Cells mediates NK Cell Activation and Parasite Control"

### Slide 1
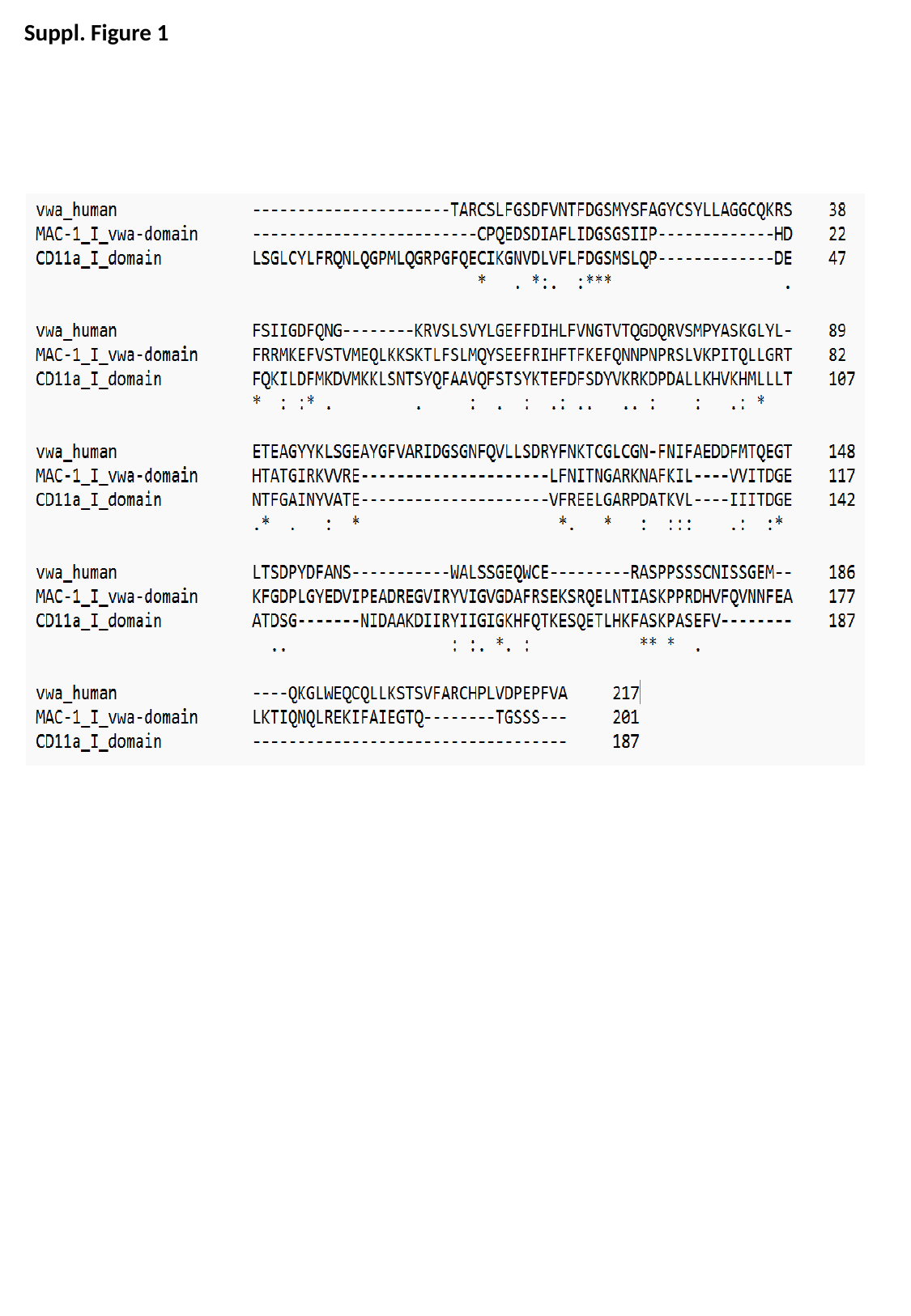

Suppl. Figure 1

### Slide 2
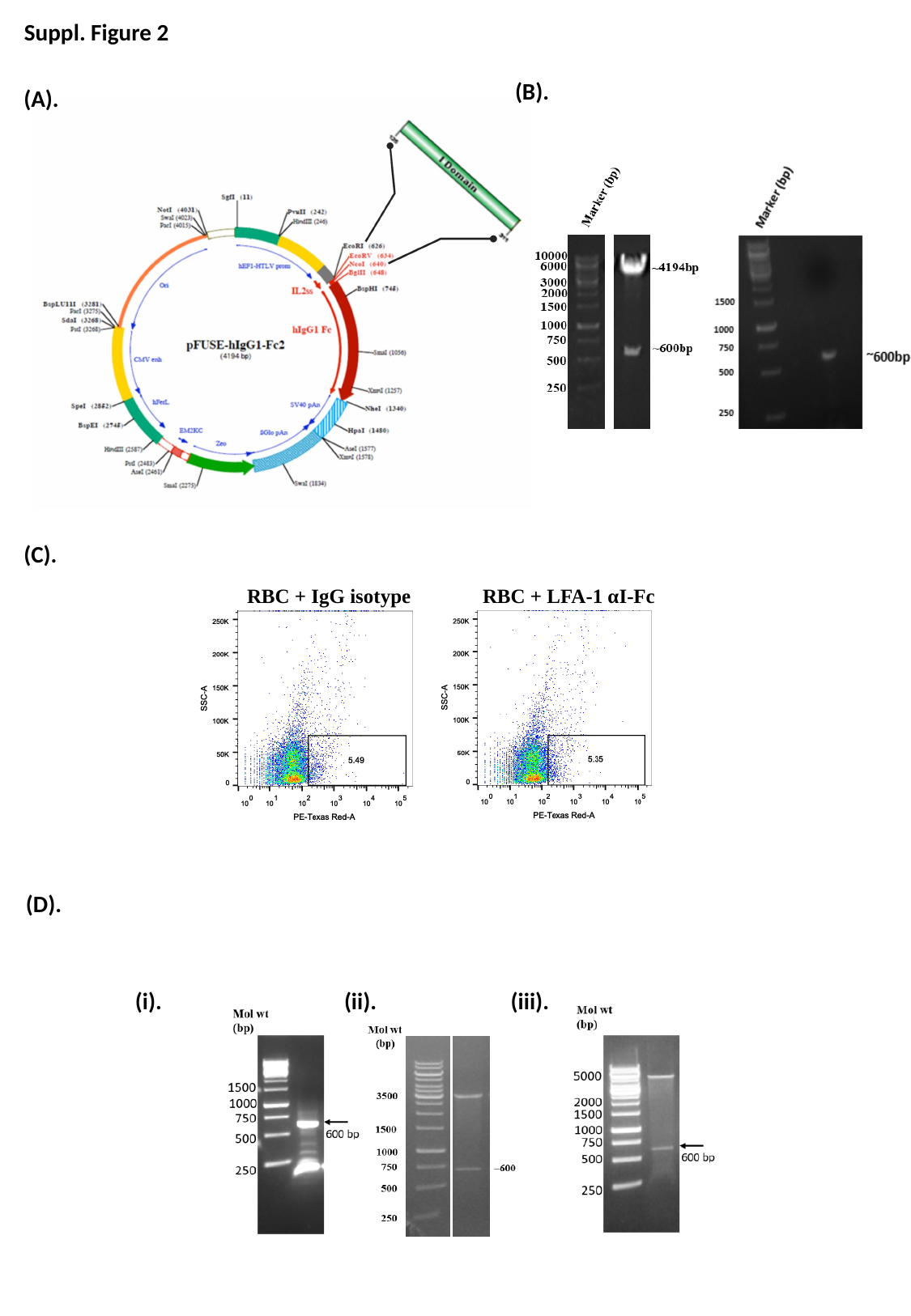

Suppl. Figure 2
(B).
(A).
(C).
RBC + IgG isotype
RBC + LFA-1 αI-Fc
(D).
(ii).
(iii).
(i).

### Slide 3
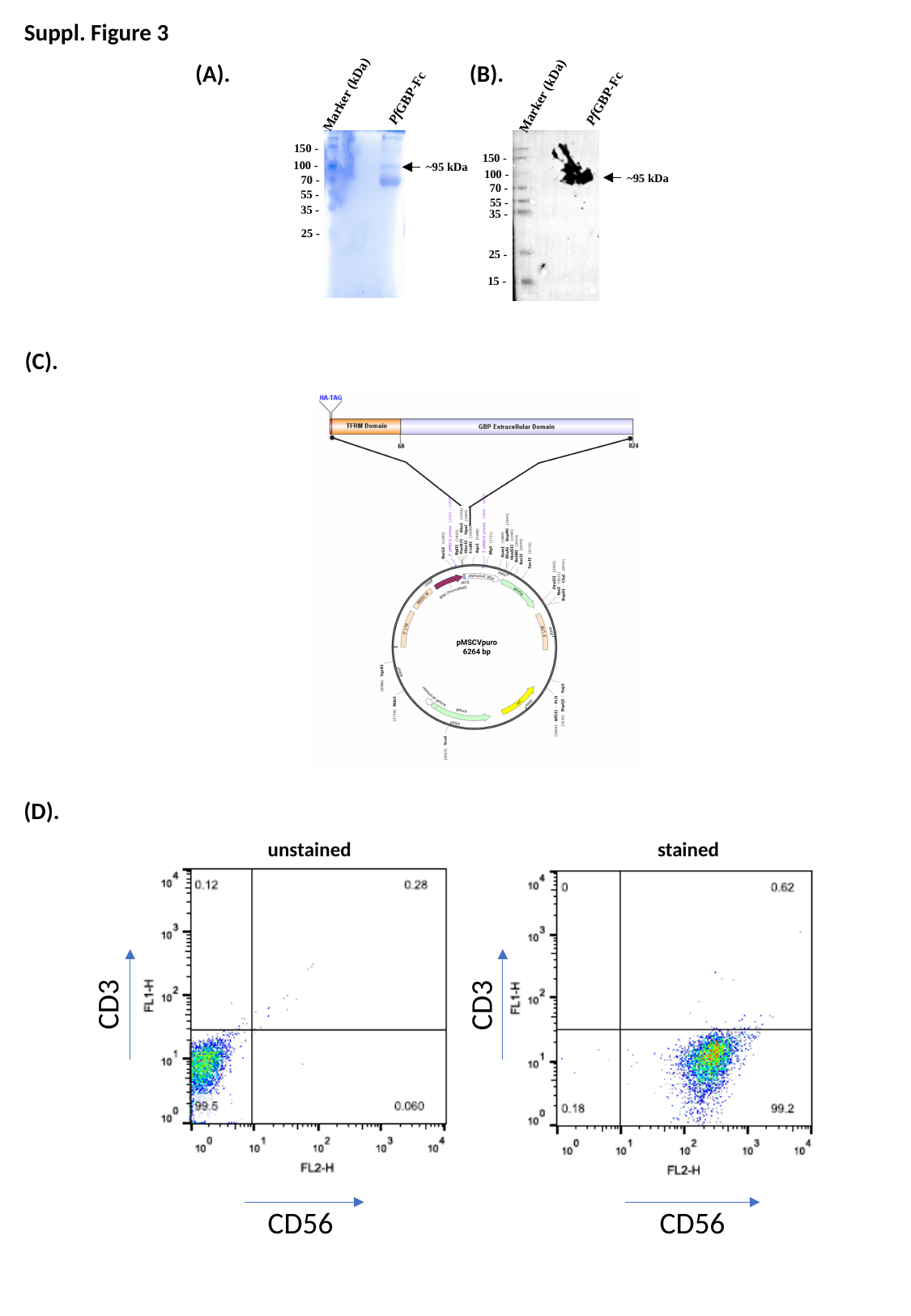

Suppl. Figure 3
(B).
(A).
Marker (kDa)
Marker (kDa)
PfGBP-Fc
PfGBP-Fc
 150 -
 150 -
 100 -
~95 kDa
 100 -
~95 kDa
 70 -
 70 -
55 -
55 -
 35 -
 35 -
25 -
25 -
15 -
(C).
(D).
unstained
stained
CD3
CD3
CD56
CD56

### Slide 4
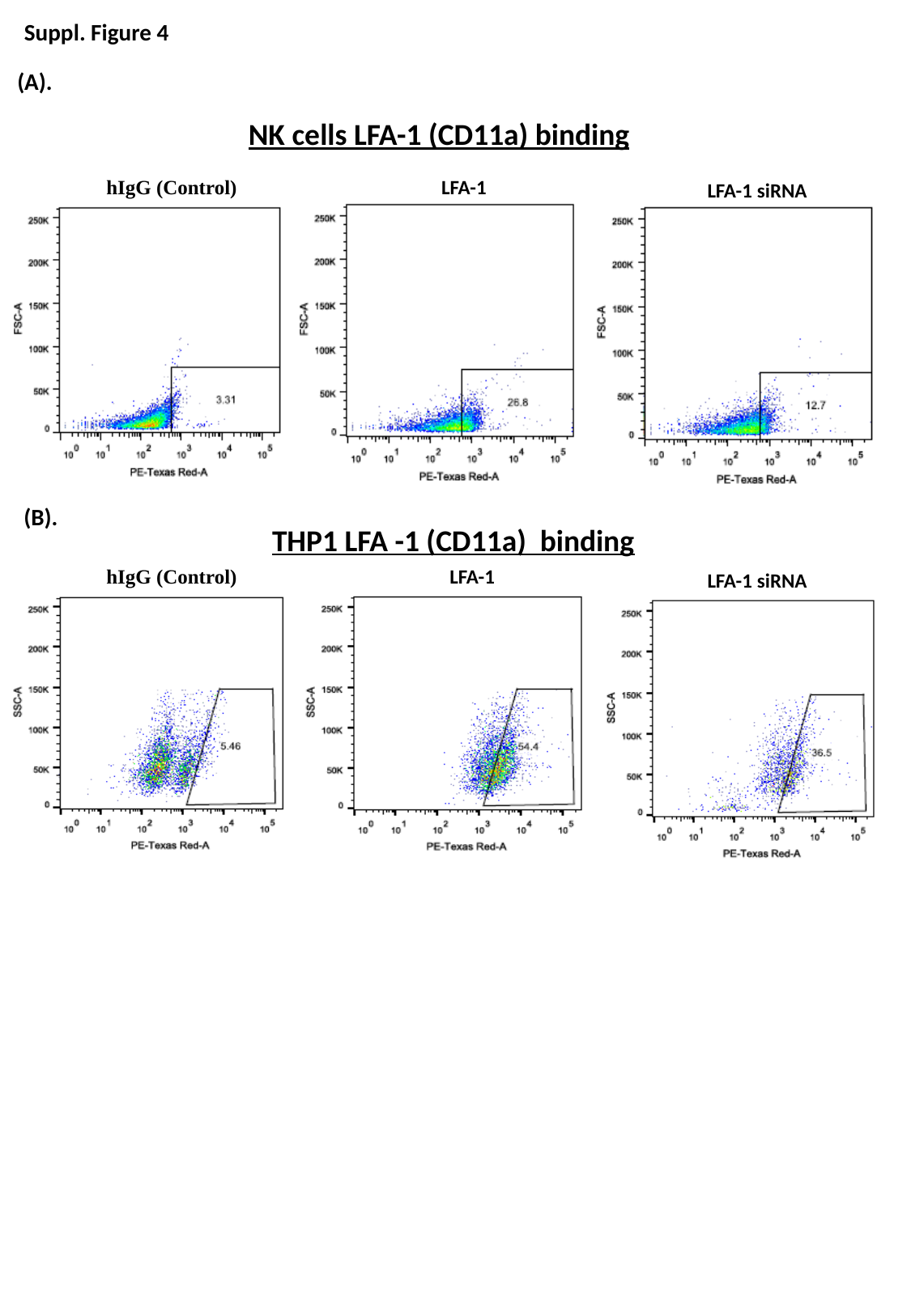

Suppl. Figure 4
(A).
NK cells LFA-1 (CD11a) binding
LFA-1
hIgG (Control)
LFA-1 siRNA
THP1 LFA -1 (CD11a) binding
LFA-1
hIgG (Control)
LFA-1 siRNA
(B).

### Slide 5
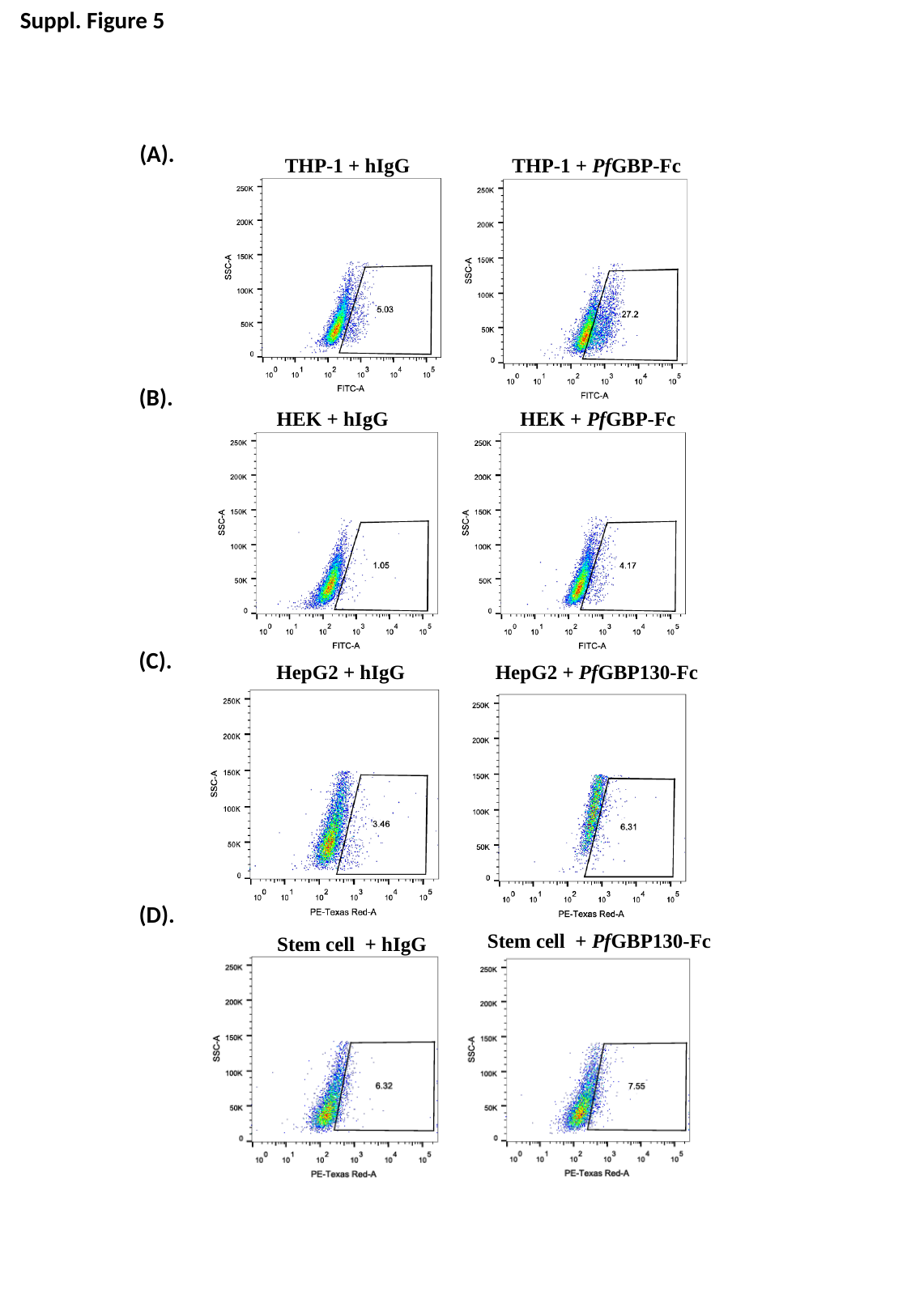

Suppl. Figure 5
(A).
THP-1 + hIgG
THP-1 + PfGBP-Fc
(B).
HEK + hIgG
HEK + PfGBP-Fc
(C).
HepG2 + hIgG
HepG2 + PfGBP130-Fc
(D).
Stem cell + PfGBP130-Fc
Stem cell + hIgG

### Slide 6
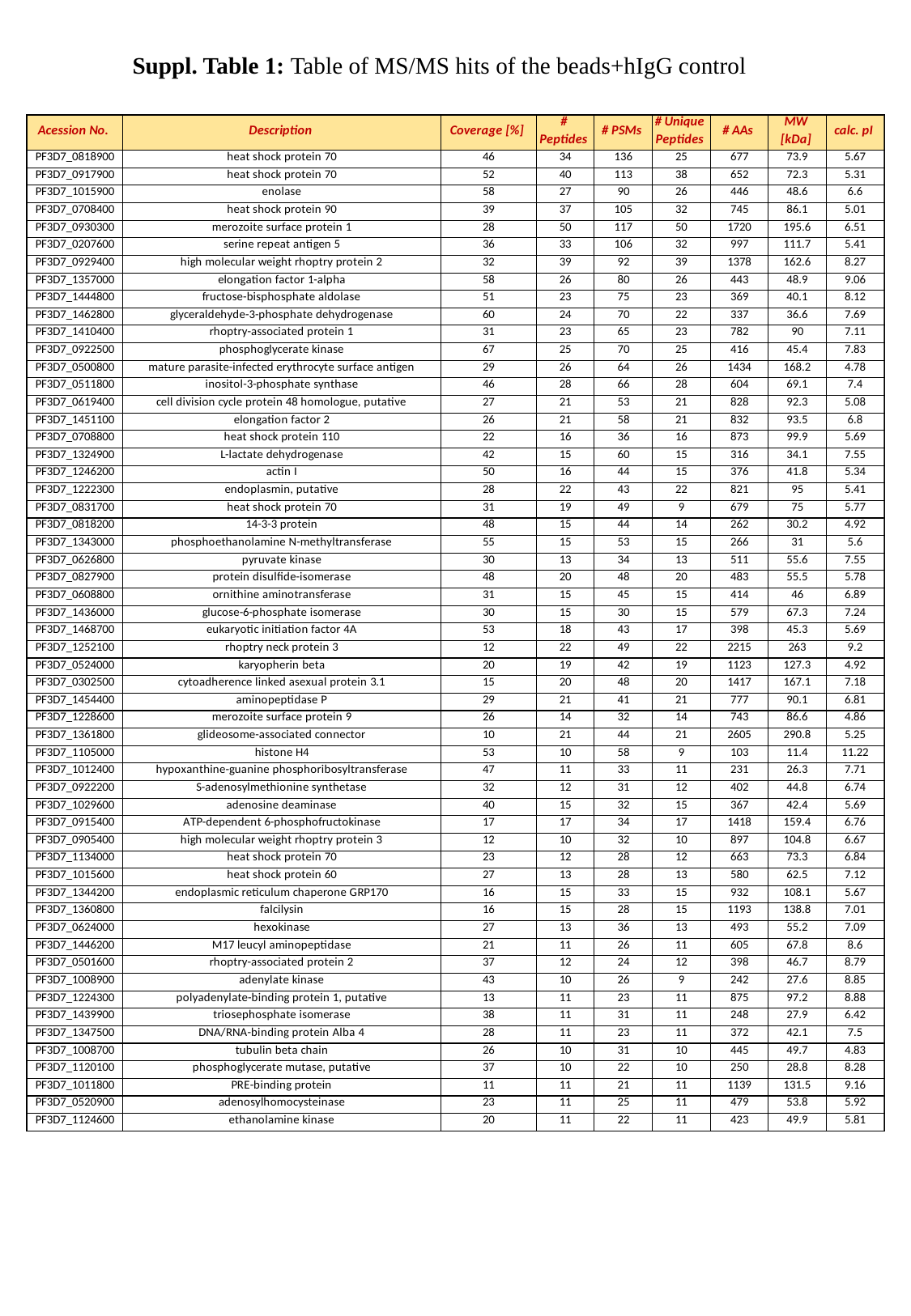

Suppl. Table 1: Table of MS/MS hits of the beads+hIgG control

### Slide 7
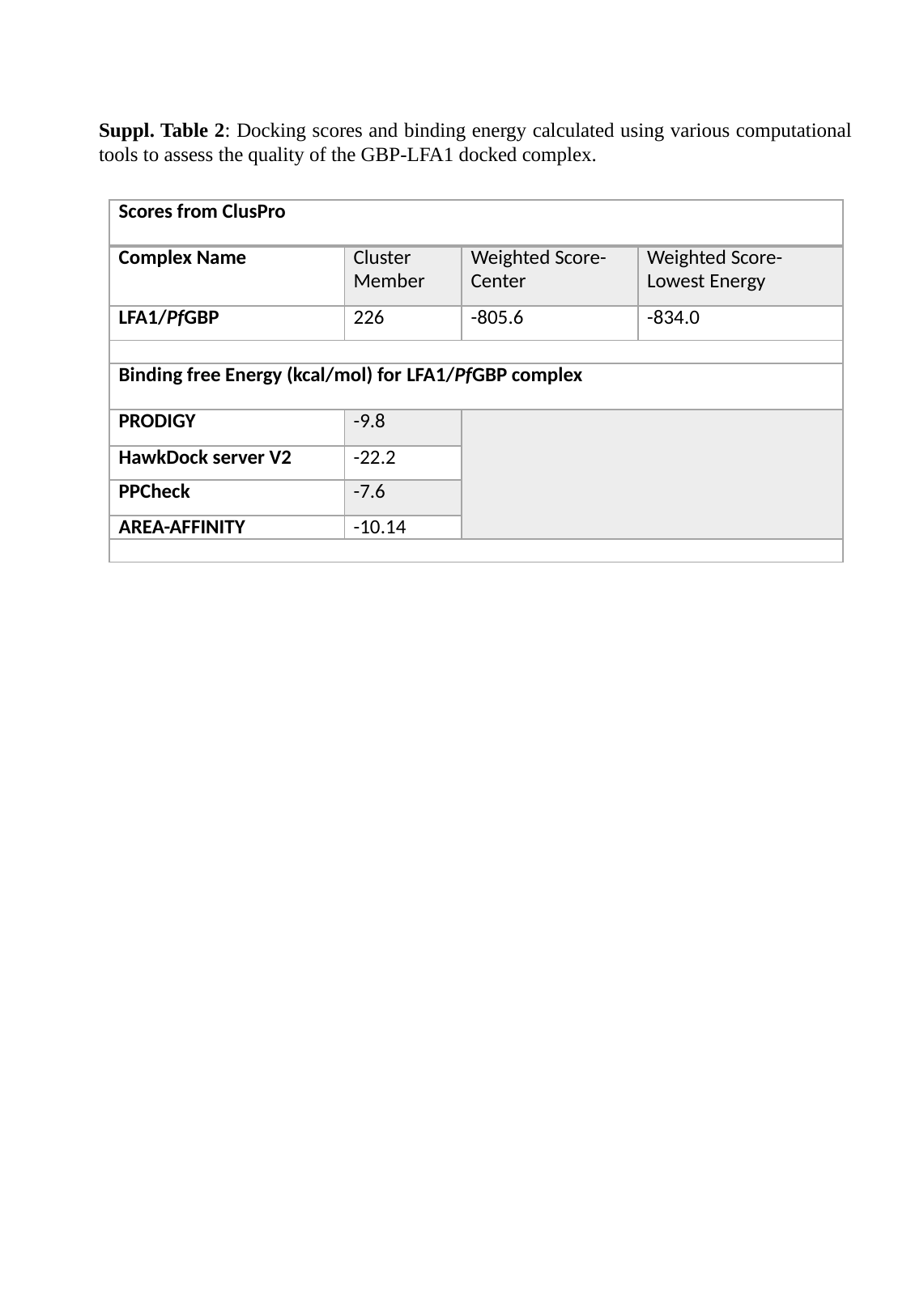

Suppl. Table 2: Docking scores and binding energy calculated using various computational tools to assess the quality of the GBP-LFA1 docked complex.
| Scores from ClusPro | | | |
| --- | --- | --- | --- |
| Complex Name | Cluster Member | Weighted Score-Center | Weighted Score-Lowest Energy |
| LFA1/PfGBP | 226 | -805.6 | -834.0 |
| Binding free Energy (kcal/mol) for LFA1/PfGBP complex | | | |
| PRODIGY | -9.8 | | |
| HawkDock server V2 | -22.2 | | |
| PPCheck | -7.6 | | |
| AREA-AFFINITY | -10.14 | | |
